# Polycomb-mediated Genome Architecture Enables Long-range Spreading of H3K27 methylation

**DOI:** 10.1101/2020.07.27.223438

**Authors:** Katerina Kraft, Kathryn E. Yost, Sedona Murphy, Andreas Magg, Yicheng Long, M.Ryan Corces, Jeffrey M. Granja, Stefan Mundlos, Thomas R. Cech, Alistair Boettiger, Howard Y. Chang

**Author notes:** These authors contributed equally to this work (Katerina Kraft, Kathryn E. Yost). These authors contributed equally to this work (Sedona Murphy, Andreas Magg).

## Abstract

Polycomb-group proteins play critical roles in gene silencing through the deposition of histone H3 lysine 27 trimethylation (H3K27me3) and chromatin compaction^1-5^. This process is essential for embryonic stem cell (ESCs) pluripotency, differentiation, and development. Polycomb repressive complex 2 (PRC2) can both read and write H3K27me3, enabling progressive spread of H3K27me3 on the linear genome^6^. Long-range Polycomb-associated DNA contacts have also been described, but their regulation and role in gene silencing remains unclear^7-10^. Here, we apply H3K27me3 HiChIP^11-13^, a protein-directed chromosome conformation method, and optical reconstruction of chromatin architecture^14^ to profile long-range Polycomb-associated DNA loops that span tens to hundreds of megabases across multiple topological associated domains in mouse ESCs and human induced pluripotent stem cells^7-10^. We find that H3K27me3 loop anchors are enriched for Polycomb nucleation points and coincide with key developmental genes, such as *Hmx1, Wnt6* and *Hoxa*. Genetic deletion of H3K27me3 loop anchors revealed a coupling of Polycomb-associated genome architecture and H3K27me3 deposition evidenced by disruption of spatial contact between distant loci and altered H3K27me3 *in cis*, both locally and megabases away on the same chromosome. Further, we find that global alterations in PRC2 occupancy resulting from an EZH2 mutant^15^ selectively deficient in RNA binding is accompanied by loss of Polycomb-associated DNA looping. Together, these results suggest PRC2 acts as a “genomic wormhole”, using RNA binding to enhance long range chromosome folding and H3K27me3 spreading. Additionally, developmental gene loci have novel roles in Polycomb spreading, emerging as important architectural elements of the epigenome.

## INTRODUCTION

Regulation of gene expression is crucial for a myriad of biological processes, including embryonic development, tissue homeostasis, and dosage compensation^2,9,10,16-18^. Gene silencing mediated by Polycomb Repressive Complexes 1 and 2 (PRC1 and PRC2) allows for temporal and tissue-specific control of gene expression during development, with aberrant regulation leading to cancer and congenital disorders^19,20^. Pluripotent stem cells, including both mouse embryonic stem cells (mESC) and human induced pluripotent stem cells (iPSC), utilize the repressive H3K27me3 mark deposited by PRC2 to suppress cell type specific expression programs, maintaining pluripotency and priming stem cells for differentiation into many lineages.

Polycomb-group proteins (PcG) and their target genes are evolutionarily conserved^21^, but how Polycomb is recruited to target genes is not fully understood. Polycomb response elements (PREs), a complex DNA element, provide a major mechanism in *Drosophila melanogaster*. In mammalian genomes, hypomethylated CG bases known as CpG islands may function similarly to *Drosophila* PREs^22,23^ and there is some evidence that Polycomb recruitment and spreading occur within 3D genome structures in addition to local spreading of H3K27me3^7-10^. However, the relationship between PcG proteins, 3D chromatin landscape and silencing of gene expression remain unclear. Several studies have revealed that PRC1 and PRC2 establish long-range interactions in various cell types and organisms^4,7,10,24-28^. Polycomb-associated interactions occur in the larger context of genome architecture and chromatin modifications, including DNA methylation^8,29^ and cohesin-dependent folding^9^, but there is emerging evidence that Polycomb can aggregate in a cohesin-independent manner^30^. In addition to DNA and chromatin-mediated recruitment, Polycomb binding to both coding and non-coding RNA has been shown to be vital to PcG recruitment^31-33^. For example, disruption of RNA-binding to EZH2, the methyltransferase subunit of PRC2, alters PRC2 recruitment to chromatin in iPSCs resulting in reduced H3K27 trimethylation on its target genes^15^. As PcGs interact with chromatin to induce compaction and mediate long-range interactions of silenced genomic regions, disruption of PRC2 recruitment via loss of RNA binding to EZH2 may alter 3D genome architecture. However, the role of PRC2 recruitment in the establishment and maintenance of chromatin architecture, in particular long-range genomic contacts^29^, and the relation to H3K27me3 propagation and gene silencing is not well known. Here we apply H3K27me3 HiChIP and optical reconstruction of chromatin architecture (ORCA) combined with genetic perturbations of H3K27me3-associated loop anchors and EZH2 binding to investigate the mechanism by which long-range polycomb associated genomic contacts are established and their role in propagation of H3K27me3 to distant sites.

## RESULTS

We performed HiChIP^12,13^ in mESCs using antibodies against two opposing histone modifications: repressive H3K27me3 deposited by PcG or the enhancer-associated H3K27 acetylation (H3K27ac)^34^. We observed 1D signal enrichment that recapitulated publicly available H3K27me3 and H3K27ac ChIP-seq datasets^35^ (**Figure 1A**) and enrichment of 1D HiChIP signal at corresponding H3K27me3 and H3K27ac ChIP-seq peaks (**Supplementary Figure 1a**). We compared high-confidence loops called by HICCUPS^36^ to those called by FitHiChIP, which additionally models the non-uniform coverage resulting from HiChIP enrichment. Over 90% of high-confidence HICCUPS loops were also detected by FitHiChIP (**Supplementary Figure 1b**) and shared loops represented a higher confidence subset of loops called by FitHiChIP (**Supplementary Figure 1c,d**). Therefore, we focused our analysis on 4,101 high-confidence H3K27me3-associated loops that were robustly detected over background and present on all chromosomes (**Supplementary Figure 1e,f**). Comparison of chromatin loops revealed that H3K27me3-associated loops bridge genomic distances spanning dozens of megabases, crossing significantly greater distances than H3K27ac-associated loops which are enriched at enhancer-promoter contact regions (p < 2.22×10^−16^, Wilcoxon rank sum test, **Figure 1A, B**). Compared to enhancer loops marked by H2K37ac, the length distribution of H3K27me3-associated loops is asymmetric and has a long tail, in particular for loops that span over 1 Mb (**Figure 1B**). The top percentile of distances is 3.33 Mb between H3K27me3-associated loop anchors versus 1.33 Mb between H3K27ac-associated loop anchors (median distance 7.9 Mb between H3K27me3-associated loop anchors for top percentile distances versus 2.0 Mb between H3K27ac-associated loop anchors).

**Figure 1:**
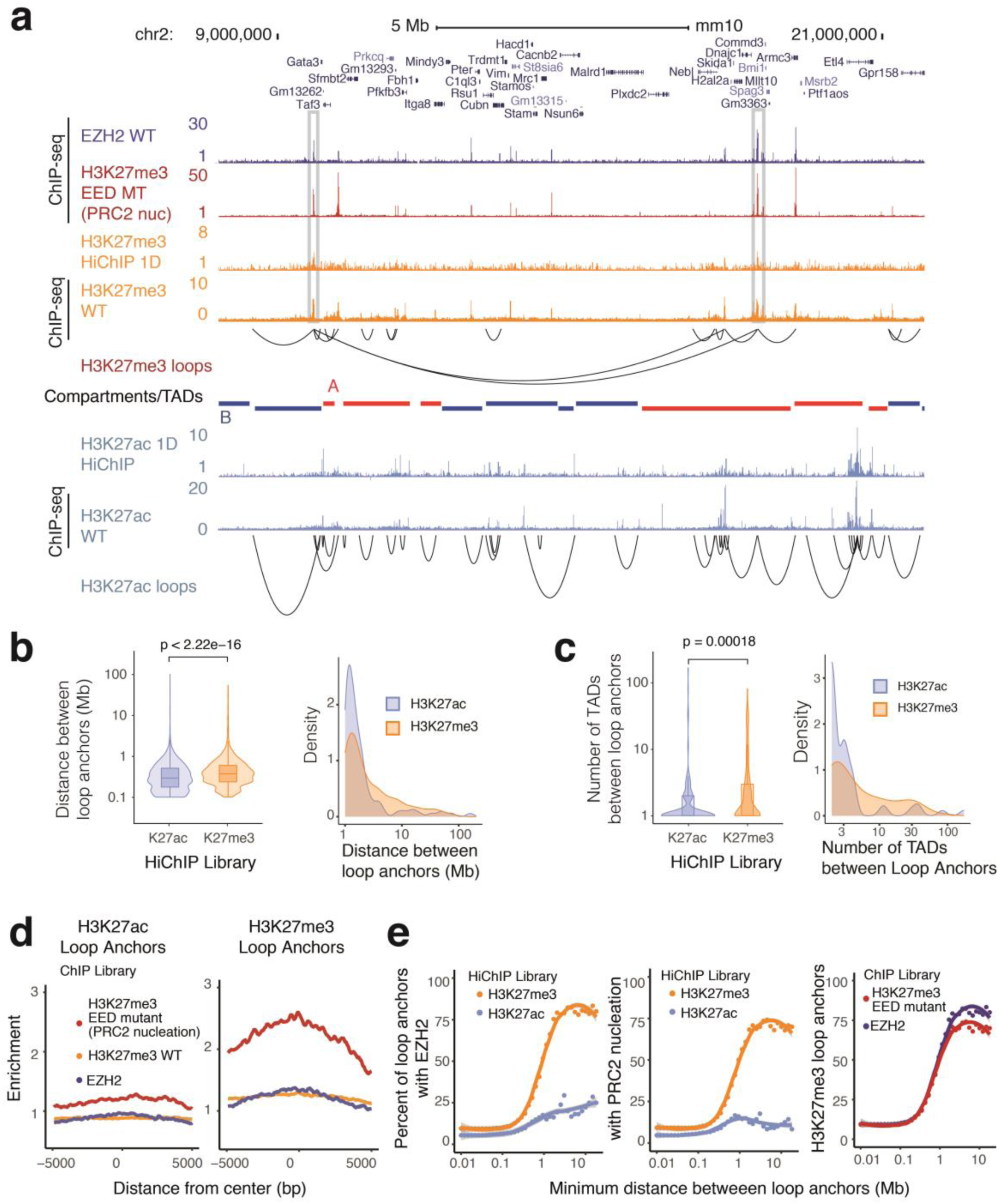
HiChIP identifies long-range polycomb-associated interactions in mESCs. **a**, HiChIP 1D signal enrichment and high-confidence HICCUPS loops for H3K27me3 and H3K27ac HiChIP at the *Gata3* locus (n = 1). H3K27me3, H3K27ac, and EZH2 ChIP-seq signal shown for comparison. Topolocigally associated domains (TADs) from mESC Hi-C data colored by A/B compartment status shown below. Selected regions with enrichment of H3K27me3 1D signal at long-range loops are highlighted. **b and c**, Density and violin plots for the **(b)** distance between H3K27me3- and H3K27ac-associated loop anchors and **(c)** number of TADs between loop anchors. P-values calculated using Wilcoxon rank sum test. **d**, Signal enrichment for EZH2, H3K27me3 WT and EED MT ChIP-seq within a 10 kb window centered on H3K27ac or H3K27me3 HiChIP loop anchors, respectively. Units of enrichment calculated as normalized ChIP-seq library depth per basepair per loop anchor. **e**, Scatter plots illustrating the relation of minimum distance between loop anchors and percentage of loop anchors overlapping EZH2 ChIP-seq peaks or EED MT H3K27me3 ChIP-seq peaks. Trend line represents less smoothed fit with span 0.6. Shaded area represents 95% confidence interval.

As enhancer-promoter contacts tend to occur specifically within topologically associating domains or TADs^26,37,38^, we next examined how H3K27me3 loops behave in regard to previously known genome architecture units, TADs bound by CTCF or larger scale chromosome A/B compartments^36^. We find that Polycomb loops can cross significantly more TADs than enhancer loops (p = 0.00018, Wilcoxon rank sum test, **Figure 1C**). The top percentile of TADs crossed is 3.6 TADs for H3K27me3-associated loops versus 2 TADs for H3K27ac-associated loops (median 10.5 TADs between H3K27me3-associated loop anchors for top percentile of loops versus 4 TADs between H3K27ac-associated loop anchors). These results suggest that long-range H3K27me3 loops may be independent units of genome architecture and are consistent with the observation that three-dimensional PcG genome architecture domains behave differently than CTCF/cohesin mediated domains^39^. Zhang et al^40^ and Boyle et al.^41^ recently identified Polycomb associated long-range DNA loops by overlapping ChIP-seq of PRC2 or PRC1 marks respectively with HiC data acquired separately, which likely represent related phenomena. The addition of H3K27me3 HiChIP data in this work demonstrates that (i) H3K27me3 and the chromosome loops are present on the same chromatin fiber; (ii) Polycomb loops can be efficiently detected at ∼1/10^th^ the sequencing cost compared to HiC (Supplementary Figure 2).

Next, we sought to identify features defining Polycomb loop anchors, specifically protein occupancy or histone modifications that are enriched at anchor points. The PcG complex PRC2 consists of core subunits: histone methyltransferase EZH2, H3K27me3 binding protein EED, architectural subunit SUZ12, and histone binding protein RBBP4^2^. Previous studies have demonstrated that mutations in the cage ring of EED disrupt interactions with H3K27me3 and lead to defects in Polycomb spreading, leading to H3K27me3 restriction at strong PcG binding-sites termed PRC2 nucleation points from which the H3K27me3 mark spreads^6^. We overlaid loop anchor points with H3K27me3 ChIP-seq from EED cage mutant mESCs (PRC2 nucleation points)^6^ and published EZH2 ChIP-seq data^42^ and found enrichment of both Polycomb nucleation points and EZH2 occupancy over H3K27me3 signal at H3K27me3 loop anchors, in contrast to H3K27ac loop anchors and all H3K27me3 peaks including non-looping peaks (**Figure 1D, Supplementary Figure 1g**). Interestingly, we find that enrichment of EZH2 occupancy and nucleation points at H3K27me3 loop anchors increases with increasing loop distance and is substantially enriched compared to H3K27ac-associated loops (**Figure 1E**). These data suggest that PRC2 complexes, especially those occupying nucleation points, may bring together distant genomic regions to establish long-range H3K27me3-associated chromatin loops capable of spanning multiple TADs and compartments.

To determine which genes are involved in H3K27me3-associated loops, we performed Gene Ontology analysis for nearest genes to loop anchors and found significant enrichment of developmentally associated processes (**Figure 2A**). While many of these loops, such as those at *Hoxa1, Hmx1*, and *Wnt6*, are also detected in deeply sequenced Hi-C^43^, we observed strong enrichment for these contacts at lower sequencing depth in H3K27me3 HiChIP when comparing virtual 4C profiles (**Supplementary Figure 2**). Many key developmental genes encoding transcription factors for cell type specification and patterning, such as the *Hoxa* gene cluster, make long-range H3K27me3-associated loops with EZH2 occupancy at both anchors, but PRC2 nucleation points occasionally only at one anchor (**Figure 2A**). While we found at least one PRC2 nucleation point for 62% of H3K27me3-associated loops spanning over 1 Mb, 37% of these had a PRC2 nucleation point at only one anchor. We sought to characterize three different examples of these developmental gene-associated loops: 1) a 31 Mb long-range loop between the *Hoxa* cluster and *Vax2* with EZH2 occupancy at both anchors, but with only *Hoxa* as a PRC2 nucleation site (**Figure 2A**); 2) a previously unobserved 3.8 Mb loop^44^ connecting *Wnt6* and *Pax3* across the *Epha4* TAD with EZH2 occupancy and PRC2 nucleation points at both anchors (**Supplementary Figure 3**); and 3) complex 2.4 Mb and 1.8 Mb loops connecting *Nkx1-1, Hmx1*, and *Msx1* with EZH2 occupancy and nucleation points at all anchor points (**Supplementary Figure 4**).

**Figure 2:**
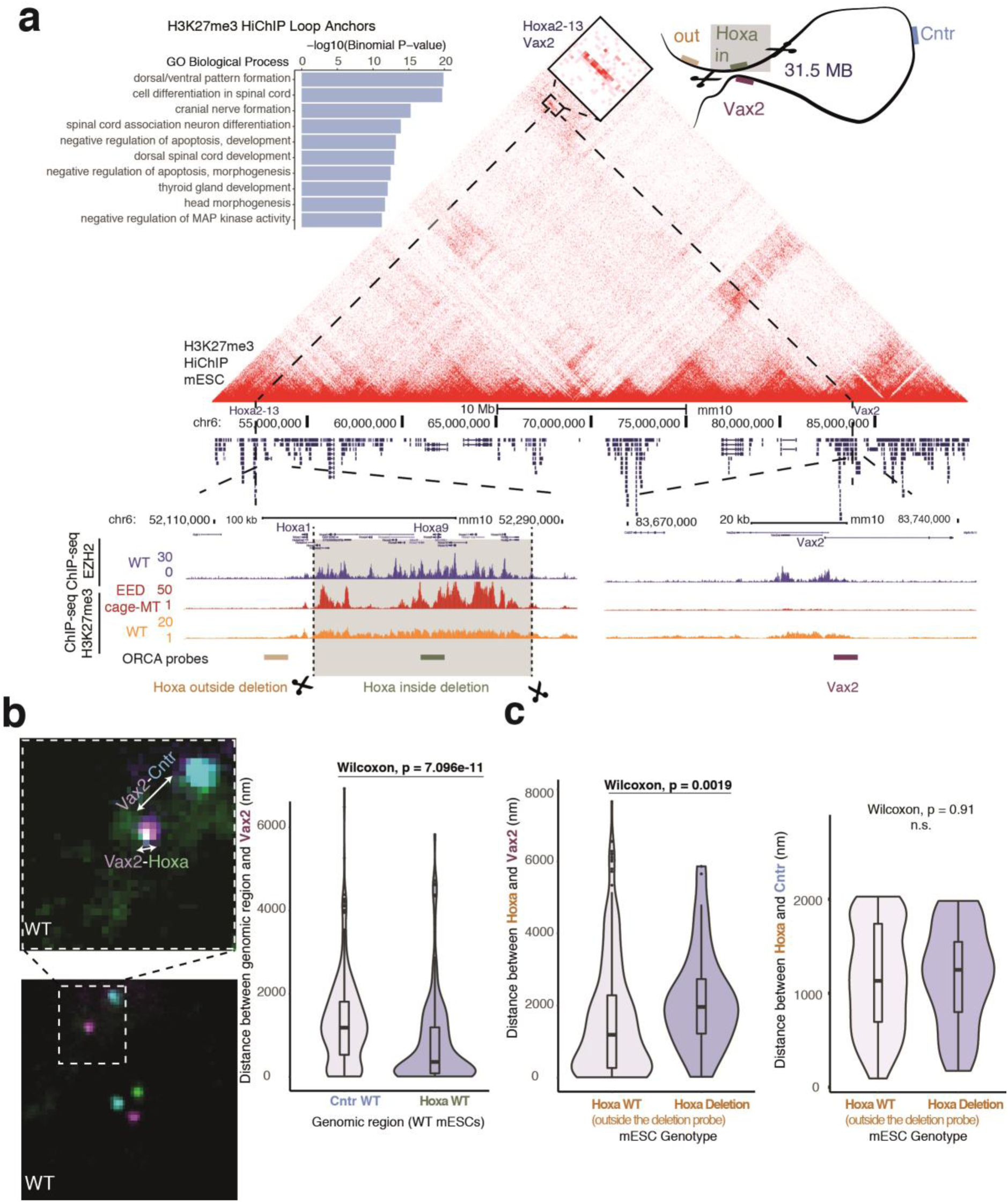
Deletion of H3K37me3-associated loop anchor at the *Hoxa* cluster alters long-range 3D interactions. **a**, Gene ontology terms enriched at mESC H3K27me3-associated loop anchors (left). H3K27me3 HiChIP contact matrix (10 kb resolution) visualizing polycomb-associated interactions at the 40 Mb region encompassing the *Hoxa* cluster and *Vax2* in mESCs. ChIP-seq signal for WT EZH2, H3K27me3 (WT and EED cage MT) and position of ORCA (<u>O</u>ptical <u>R</u>econstruction of <u>C</u>hromatin <u>A</u>rchitecture) probes is shown below. **b**, ORCA imaging of three probes targeting *Hoxa, Vax2*, and Control (Cntr) regions demonstrates interaction between *Hoxa* cluster with *Vax2* at the single-nucleus level in wild-type mESCs. Violin plots of the distance between *Vax2* and the *Hoxa* or *Cntr* probes (WT cells n = 2190; MT cells n = 520). *Hoxa* probe located within the polycomb-associated loop anchor used in wild-type mESCs. **c**, Violin plots of the distance between *Hoxa* and *Vax2* (as measured by ORCA) for WT and Hoxa Deletion mESCs (left). Violin plots of the distance between *Hoxa* and *Cntr* (as measured by ORCA) for WT and Hoxa Deletion mESCs (right). As the *Hoxa* polycomb-associated loop anchor is deleted in Hoxa deletion mESCs, a probe targeting a region adjacent to *Hoxa* loop anchor was used for both wild-type and Hoxa deletion mESCs. Wilcoxon test for significance.

To test the relationship between long-range looping and H3K27me3 spreading by PRC2 we used CRISPR-Cas9 editing to generate homozygous deletion alleles of loop anchors containing both PRC2 nucleation and EZH2 occupancy sites (**Figure 2A**). To interrogate the effects of these deletions and provide an orthogonal measurement of long-range H3K27me3-associated contacts, we monitored resulting genome architecture changes using ORCA (Optical Reconstruction of Chromatin Architecture), combining multiplexed DNA FISH to sequentially image DNA loci at resolutions higher than the diffraction limit^14^ (**Figure 2B**). To enable ORCA following CRISPR-Cas9 editing, we designed probes to target both the original loop anchor, such as the *Hoxa* cluster, as well as a region adjacent to the deletion. Additional probes were placed at the *Vax2* loop anchor 31 Mb away and at a control region located at the midpoint of the two loop anchors in linear genomic distance. In order to fully remove the PRC2 nucleation and EZH2 occupancy site, we deleted the majority of the *Hoxa* cluster from *Hoxa2* to *Hoxa13*, leaving *Hoxa1* which is outside of loop anchor detected by HiChIP. *Hoxa* cluster genes *Hoxa2* to *Hoxa13* included in the deletion are not expressed in mESC so the effects of deletion should not be attributed to altered dosage of deleted genes. ORCA confirmed the presence of a 31 Mb loop identified by HiChIP in unedited cells, with 40% of cells positive for contact (<150 nm) between *Hoxa* and *Vax2* by high-resolution imaging (**Figure 2B**). In cells where the *Hoxa* loop anchor was deleted there was a significant increase in the physical distance between *Vax2* and *Hoxa* (**Figure 2C**). This supports the importance of the loop anchor which contains EZH2 occupancy and PRC2 nucleation points in establishing long-range contacts, which upon removal affects the spatial organization of regions outside of the original loop anchor.

We next asked if deletion of H3K27me3 loop anchors can affect spreading of the Polycomb-mediated H3K27me3 mark. We performed H3K27me3 Cut&Tag^45^ in mESCs with loop anchor deletions for each anchor deletion at *Hoxa20-13, Wnt6*, or *Hmx1* and identified genomic loci with differential H3K27me3 deposition, focusing on significantly altered sites *in cis* more likely to be directly impacted by loop anchor deletion (**Figure 3A**). Because *Hoxa2-Hoxa13, Wnt6*, and *Hmx1* genes are not expressed in mESCs, the effects we observe are most likely to result from deleting the genomic region and not from eliminating expression of these genes. Our data showed reduced H3K27me3 adjacent to deletion breakpoints, consistent with loss of local Polycomb spreading (**Figure 3B**). Surprisingly, we observed many differences in H3K27me3 at distant sites tens of megabases from the deletion site, most strikingly in the *Hoxa* deletion, with unaltered H3K27me3 sites in between (**Figure 3A,C**). To investigate the cause of such distant effects, we examined the relationship between EZH2 occupancy and found that sites which lose H3K27me3 lack intrinsic EZH2 occupancy (**Figure 3C,D, Supplementary Figure 5**). These regions also tend to co-localize in the same compartment (**Figure 3E**). Interestingly, we observed no significant changes in H3K27me3 at long-range contacts with deleted loop anchors identified by HiChIP, all of which have EZH2 occupancy. This data suggests an intriguing possibility that PRC2 nucleation sites such as the *Hoxa* gene cluster may distribute the repressive H3K27me3 mark to distant regions within the same compartment which lack intrinsic Polycomb binding. We predict that these interactions are dynamic and transient, as we did not identify significant looping interactions between the deleted nucleation sites and regions with loss of H3K27me3 in our HiChIP data. We also identified sites with increased H3K27me3 at local as well as distant sites (**Figure 3C**), consistent with recent work demonstrating global redistribution of PcG complexes upon pertrubations such as BAF depeletion^46^.

**Figure 3:**
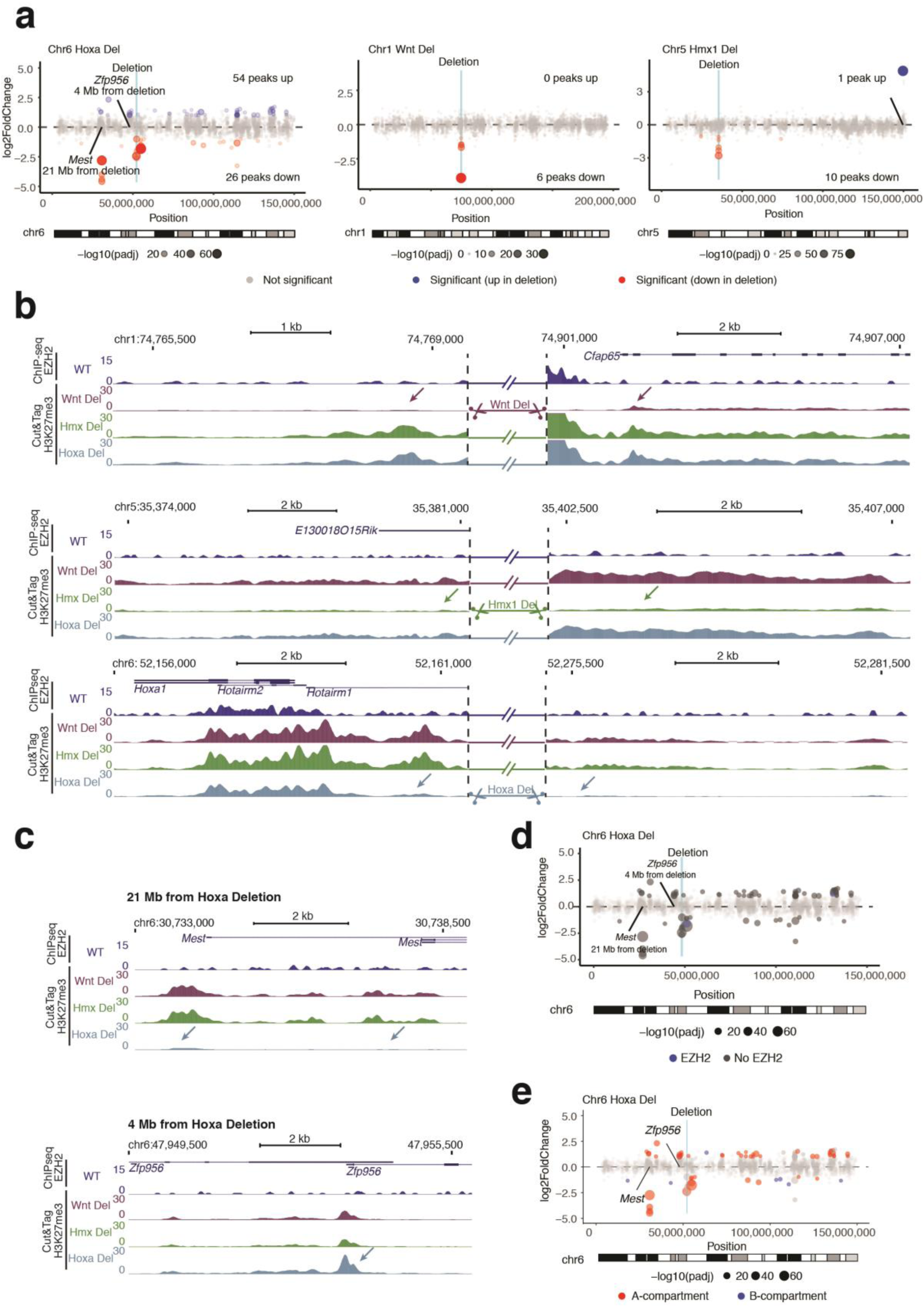
Deletion of polycomb-associated loop anchors leads to both local and long-range changes in H3K27me3 modification *in cis*. **a**, Scatter plots for the three different anchor point deletions (*Hoxa, Wnt, Hmx1*) illustrating the effects on altered H3K27me3 Cut&Tag signal versus the genomic position on the chromosome. Log2 fold changes and p-values (cutoff of absolute value log2FC > 1 and Benjamini-Hochberg adjusted p-value < 0.05 for significance) calculated in DESeq2 for each anchor point deletion clone (n=3 replicates) relative to others. **b**, Local changes in vicinity of anchor point deletions are depicted by H3K27me3 Cut&Tag signal tracks. Arrows indicate significantly altered regions in mutants. EZH2 ChIP-seq signal in WT mESCs is shown above. Signal at deleted regions omitted for clarity. **c**, Long-range alterations in H3K27me3 resulting from deletion of *Hoxa* loop anchor, resulting both in down- and upregulation (indicated with arrows). **d and e**, Scatter plots as in **(a)** but showing the relation of H3K27me3 signal depending on **(d)** the occupancy of EZH2 at the altered site or **(e)** A/B compartment status.

To determine if long-range Polycomb loops are conserved features across evolution, we performed H3K27me3 HiChIP in human induced pluripotent stem cells (iPSCs). We lifted over mESC H3K27me3-associated loops anchor coordinates to the human genome and found that approximately 30% of high confidence loops identified by H3K27me3 HiChIP in human iPSCs were shared with mouse ESCs, and these conserved H3K27me3 loops in the human genome were enriched for similar developmental Gene Ontology terms as observed in mESCs (**Supplementary Figure 6A**).

To examine the global effects of PRC2 binding on genome architecture, we utilize a recently published EZH2 variant with mutations in two regions (F32A, R34A, D36A, K39A; 489-494 PRKKKR to NAAIRS) that is deficient in RNA binding but otherwise has normal PRC2 complex formation and intact H3K27me3 methylase activity^15,47^. This RNA-binding deficient EZH2 mutant (*EZH2*^*RNA-*^ hereafter) results in global alterations in Polycomb binding and H3K27me3 deposition(**Figure 4A**)^15^, raising the question of whether RNA binding may be relevant to the long-range Polycomb spreading described here. To ask how EZH2’s promiscuous RNA binding may contribute to H3K27me3-associated chromatin architecture, we performed H3K27me3 HiChIP in wildtype and *EZH2*^*RNA-*^ iPSCs. *EZH2*^*RNA-*^ iPSCs had significant reduced contacts at Polycomb anchor sites at key developmental loci, including *HOX, PAX, NKX*, and *TBX* (**Supplementary Figure 6B**). Because *EZH2*^*RNA-*^ iPSCs have decreased H3K27me3 at these same loci, resulting changes in HiChIP signal can be attributed to either loss of H3K27me3 modification or loss of contact. Therefore, we performed 4C-seq^48^, a targeted DNA proximity assay not dependent on ChIP enrichment, at viewpoints in regions with differential contact identified by HiChIP such as an H3K27me3-associated loop that spans the adjacent *PAX9* and *NKX2-1* locus to the *FOXA1* locus ∼500 kilobases away. *PAX9-NKX2-1* shows strong EZH2 occupancy while *FOXA1* has modest EZH2 occupancy in wild type cells (**Figure 4B**). In *EZH2*^*RNA-*^ iPSCs, *PAX9-NKX2-1* no longer comes into proximity with *FOXA1* as shown by 4C with two independent viewpoints and primer sets, and concordantly *FOXA1* loses EZH2 occupancy (**Figure 4B**). Analysis of the *TBX3* and *TBX5* loci shows a similar deficit of the *EZH2*^*RNA-*^ mutant in Polycomb-mediated chromosome looping and spread of EZH2 occupancy from the putative nucleation point (*TBX3*) to its loops partner (*TBX5*), but minimal effects on chromosome looping at sites which maintain EZH2 occupancy (**Supplementary Figure 6C**). These results suggest that RNA binding by EZH2 is required to drive long-range chromosome looping and spread Polycomb occupancy to distant loci.

**Figure 4:**
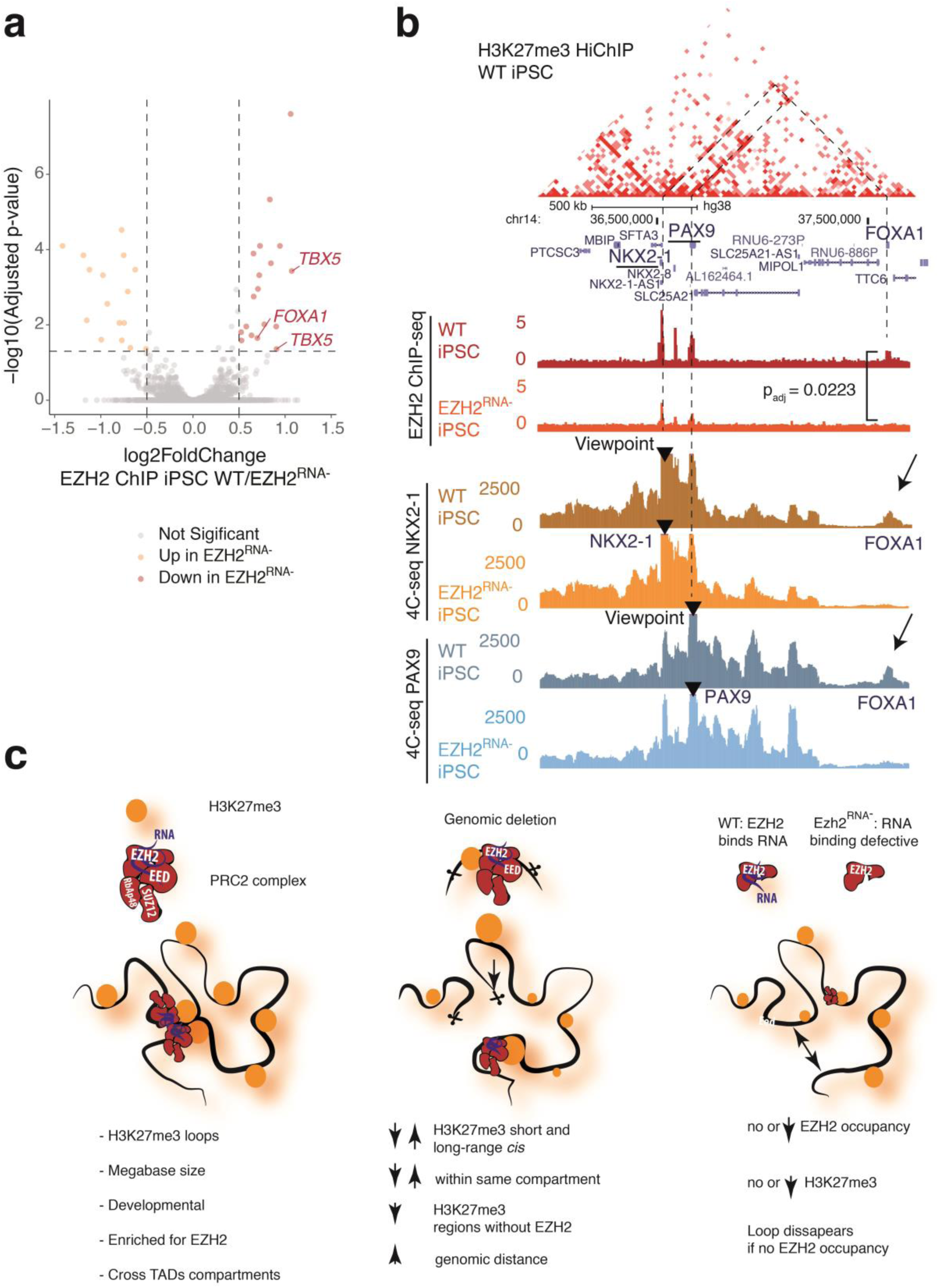
Altered polycomb binding due to loss of RNA binding by EZH2 alters genome architecture in human iPSCs. **a**, Volcano plot of differential EZH2 ChIP-seq signal (WT/*EZH2*^*RNA-*^ iPSCs) (right). Log2 fold changes and p-values (cutoff of absolute value log2FC > 0.5 and Benjamini-Hochberg adjusted p-value < 0.05 for significance) calculated in DESeq2. **b**, WT H3K27me3 HiChIP contact matrix at *NKX2-2*/*PAX9*/*FOXA1* locus with H3K27me3 and EZH2 ChIP-seq and 4C-seq at *NKX2-1* and *PAX9* viewpoints in WT and *EZH2*^*RNA-*^ iPSCs. Lost contact with *FOXA1* accompanied by loss of EZH2 binding highlighted with Benjamini-Hochberg adjusted p-value for EZH2 ChIP-seq signal (WT/*EZH2*^*RNA-*^ iPSCs) shown. **c**, Summary of the findings. In WT stem cells, polycomb-associated H3K27me3 loops connect vast genomic distances spanning dozens of megabases, crossing TADs and A/B compartments. Deletion of anchor points leads to both local and distal changes of H3K27me-spreading *in cis*, preferentially affecting regions which lack EZH2 occupancy and are located in the same compartment as the original anchor. RNA-binding deficient mutant EZH2 results in loss of looping at loci at sites with reduced EZH2 occupancy.

## DISCUSSION

Here, we apply H3K27me3 HiChIP to identify long-range Polycomb-associated loops among developmental genes, and we demonstrate that local or global loss of Polycomb binding can mediate long-range effects on H3K27me3 deposition and genome architecture. Polycomb loops link H3K27me3–modified loci from the same chromatin compartment that are separated by tens to hundreds of megabases on the linear chromosome, and they appear orthogonal to the ∼1Mb size TADs based on CTCF and cohesin, indicating an intermediary structure in the hierarchical organization of 3D genome folding. This finding is consistent with studies that conclude that PRC-1 associated interactions are independent of CTCF and TADs^9,41^. While changes in enzymatic activity of PRC1 do not impair these interactions, our study demonstrates loss of Polycomb-associted interation when disrupting PRC2 RNA binding domain, suggesting there may be distinct roles for PRC1 and PRC2 in regulation of genome architecture^41^. Zhang et al. recently reported long-range chromosomal loops in hematopoietic stem cells marked by nardis of DNA methylation and high H3K27me3^29^; thus Polycomb loops may be a conserved feature of embryonic and adult tissue stem cells.

Our results also suggest a new view of developmental gene loci as architectural elements of the epigenome, nucleating and spreading H3K27me3--a role analogous to centromeres and telomeres that mediate position effect variegation through the spread of H3K9me3^49^. Because all three of the TF loci we deleted are not transcribed in ESCs, the effects of locus deletion on long-range H3K27me3 deposition are likely due to architectural roles of these loci as noncoding regulatory DNA elements. These effects should be considered when investigators interpret large-scale deletions of *Hox* and other developmental gene loci. We find that EZH2 occupancy, specifically at sites mediated by RNA binding, is essential in establishment of long-range genomic contacts and spreading of H3K27me3, potentially connecting widely separated chromatin regions as a “genomic wormhole” (**Figure 4C)**. PRC2 binds thousands of RNAs on chromatin^32,33,50^ and inhibition of EZH2-RNA interactions results in globally reduced EZH2 occupancy, altered H3K27me3 modification, and differentiation defects^15,51^.

Many developmental TF loci encode positionally conserved noncoding RNA transcripts^52^, which may facilitate PRC2 spreading and enforcement of chromosome contacts. While the specific RNA species involved and detailed recruitment mechanisms should be addressed in future studies, the results of this study suggest that Polycomb-associated contacts may be important for proper gene regulation during development.

## Acknowledgements

We thank M. Mumbach for HiChIP protocol optimization, X. Ji, D. Wagh and J. Coller for sequencing support, and members of Chang and Boettiger labs for discussion. Supported by NIH RM1-HG007735 (to H.Y.C.), DFG KR 5172/1 (to K.K.), NIH K99GM132546 (to Y.L.). K.E.Y. was supported by the National Science Foundation Graduate Research Fellowship Program (NSF DGE-1656518) and a Stanford Graduate Fellowship. SM was supported by the DFG grant MU 880/16-1. Sequencing was performed by the Stanford Functional Genomics Facility (supported by NIH grant S10OD018220). H.Y.C. and T.R.C. are Investigators of the Howard Hughes Medical Institute.

## Author Contributions

K.K. and H.Y.C. conceived the project. K.K., K.E.Y., A.M., Se.M., and Y.L. performed experiments. K.E.Y. analyzed genomic data with input from K.K., Se.M., A.M., Y.L., M.R.C., and J.M.G. Se.M. analyzed imaging data. St.M., T.R.C., A.B. and H.Y.C. guided data analysis. K.K., K.E.Y., and H.Y.C. wrote the manuscript with input from all authors.

## Data availability

All sequencing data generated in this study are available through the Gene Expression Omnibus (GEO) under accession number GSEXXXXXX. GEO dataset will be made publicly available upon publication of peer-reviewed paper.

## SUPPLEMENTARY FIGURES

**Supplementary Figure 1:**
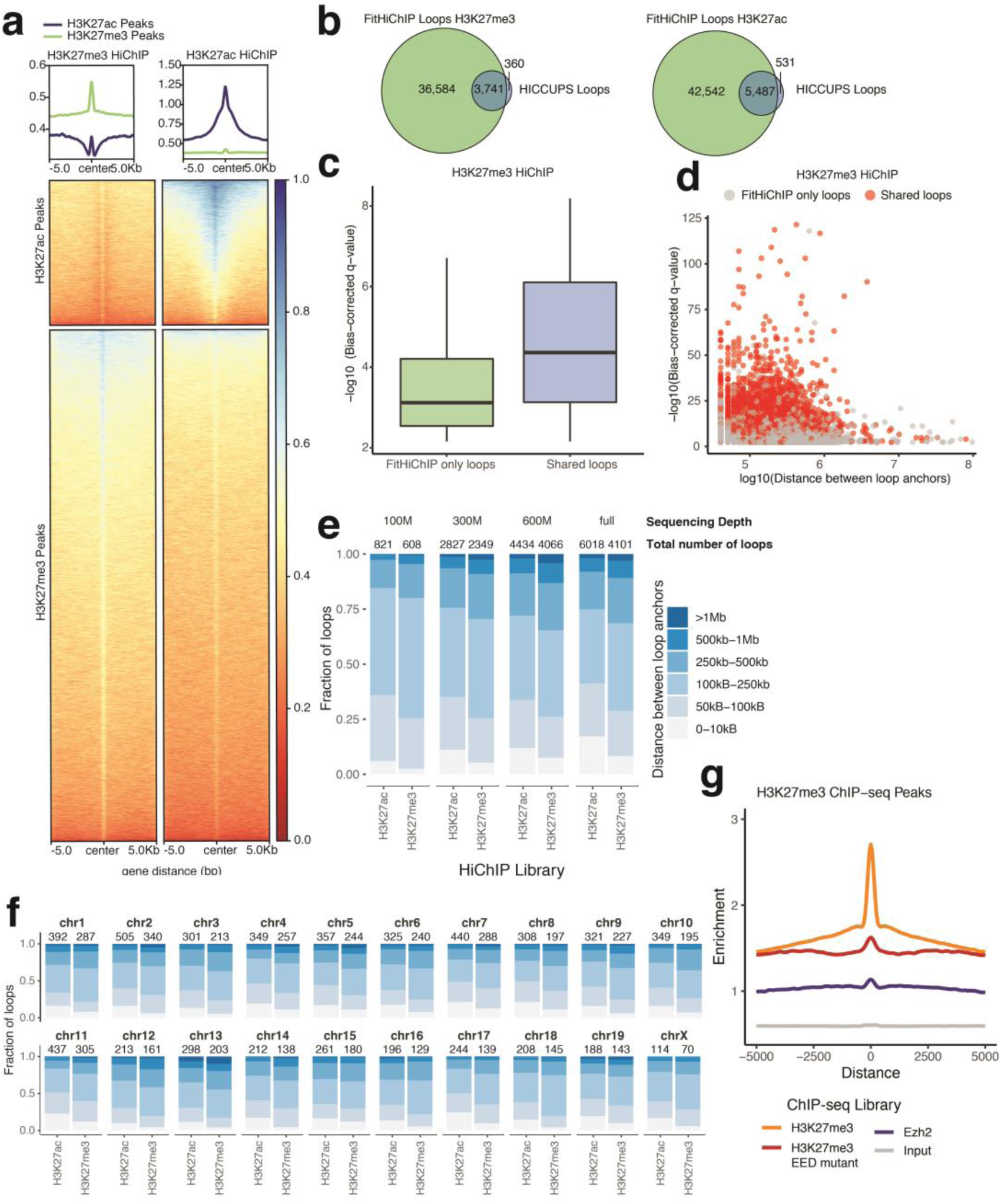
Quality control for mESC H3K27me3 and H3K27ac HiChIP. **a**, Enrichment of 1D HiChIP signal at H3K27me3 and H3K27ac peaks (ENCODE)^35^. **b**, Overlap between loops called by FitHiChIP and HICCUPS for mESC H3K27ac and H3K27me3 HiChIP. Loops were considered to be shared if both anchors overlap. **c**, Boxplot of FitHiChIP bias-corrected q-values for H3K27me3 HiChIP loops called only by FitHiChIP and those shared with loops called by HICCUPS. **d**, Scatter plot of loop distance versus bias-corrected q-value for H3K27me3 HiChIP loops called only by FitHiChIP and those shared with loops called by HICCUPS. **e**, Bar plot of called HiChIP loops separated by distance between loop anchors for H3K27ac and H3K27me3 HiChIP in mESCs. Loops were called after downsampling to 100, 300 and 600 million reads with similar results despite fewer total loops called. **f**, Bar plot of called HiChIP loops separated by distance between loop anchors and chromosome for H3K27ac and H3K27me3 HiChIP in mESCs, demonstrating precense of long-range H3K27me3 on all chromosomes. **g**, Signal enrichment for EZH2, H3K27me3 WT and EED MT ChIP-seq within a 10 kb window centered on H3K27me3 ChIP-seq peaks. Representative input control from H3K27me3 ChIP-seq included. Units of enrichment calculated as normalized ChIP-seq library depth per basepair per loop anchor.

**Supplementary Figure 2:**
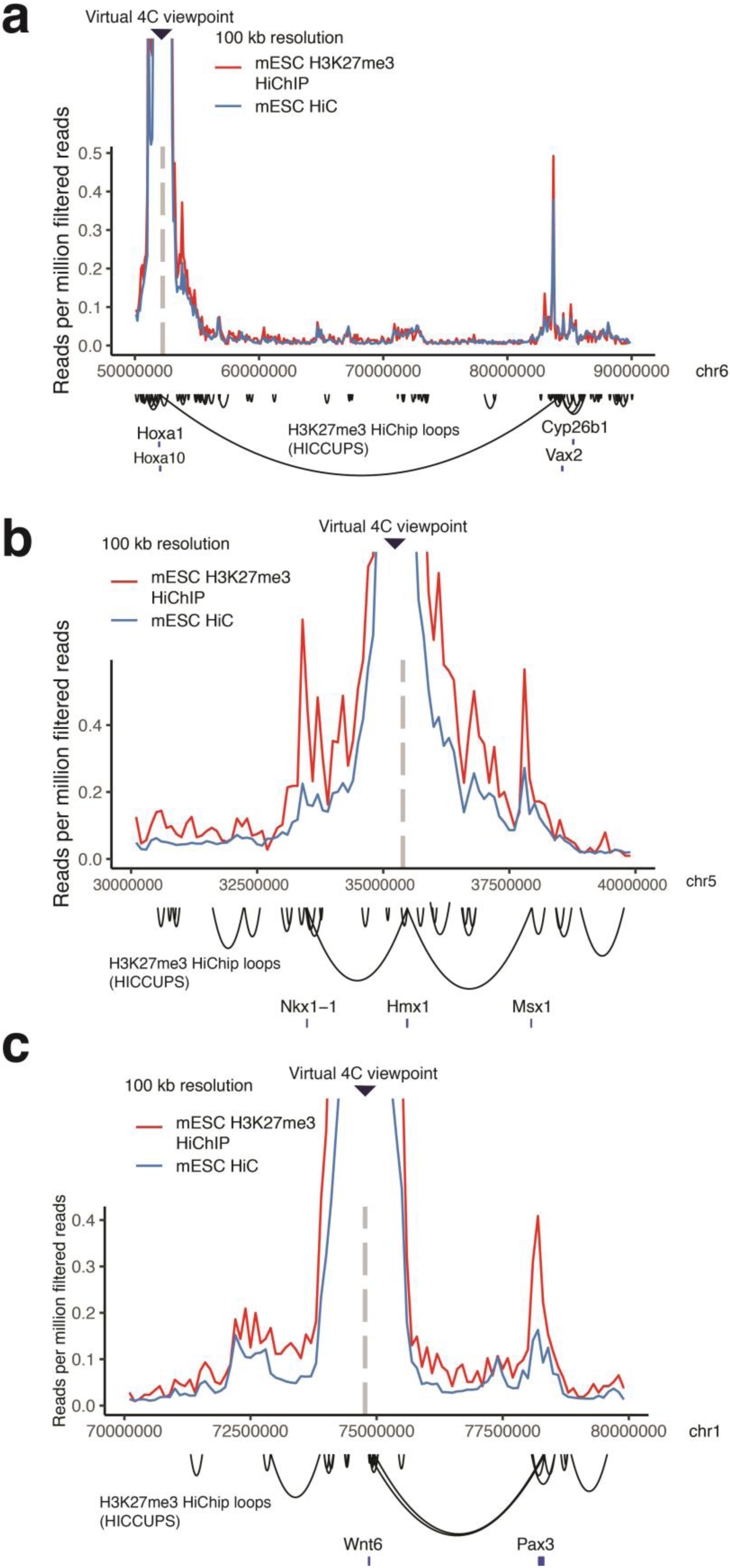
Virtual 4C comparison of mESC Hi-C and mESC H3K27me3 HiChIP. **a**, Virtual 4C interaction profile at the *Hoxa1* promoter in mESCs for Hi-C^43^ (Bonev, *et al*, 7,260,480,082 total sequencing read pairs, 4,255,891,410 filtered read pairs) and H3K27me3 HiChIP (this study, 644,288,043 total sequencing read pairs, 215,356,878 filtered read pairs) scaled by number of filtered read pairs. **b**, Virtual 4C interaction profile at the *Hmx1* promoter. **c**, Virtual 4C interaction profile at the *Wnt6* promoter.

**Supplementary Figure 3:**
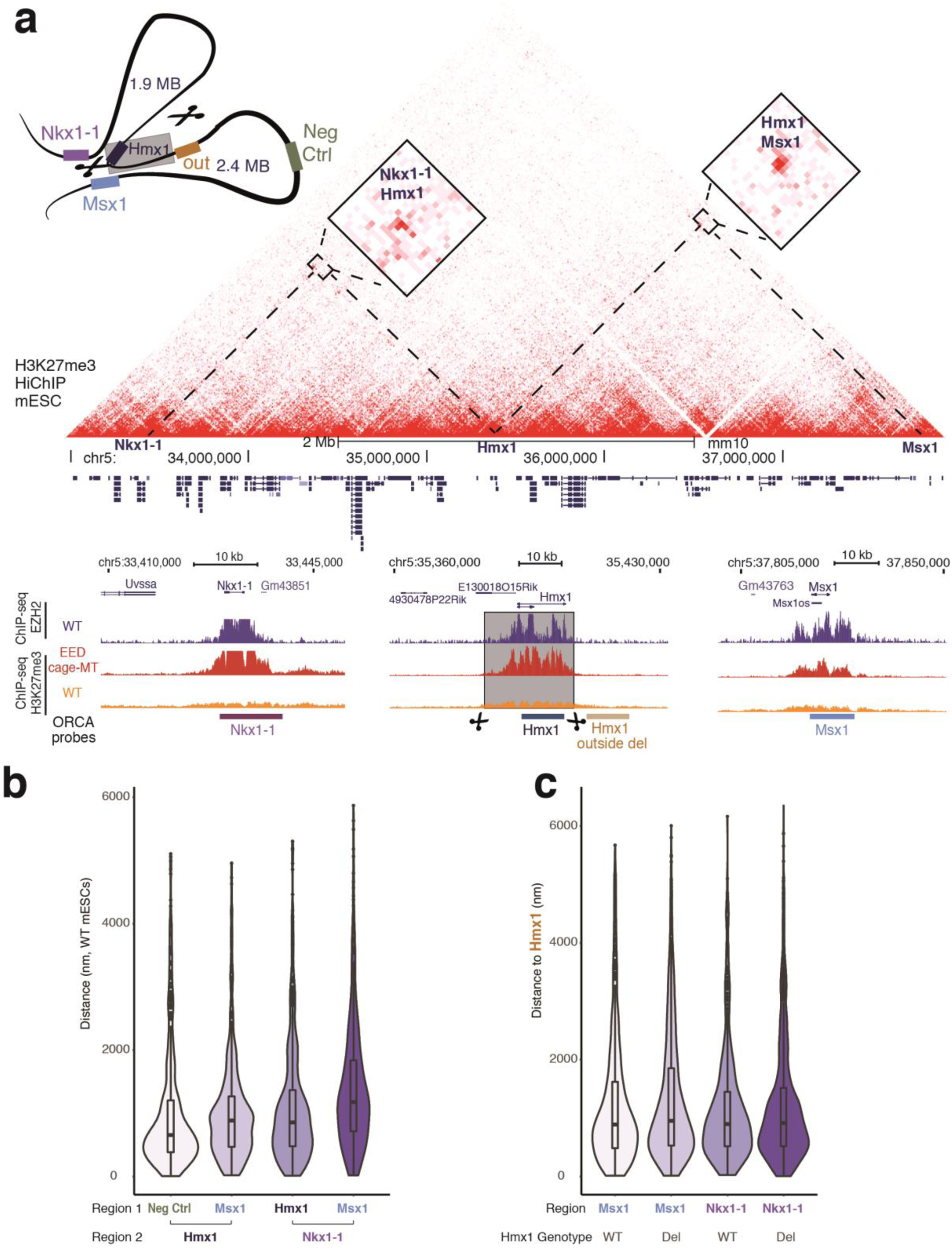
Hmx1 loop anchor deletion. **a**, H3K27me3 HiChIP contact matrix (10 kb resolution) at the *Nkx1-1*/*Hmx1*/*Msx1* locus in WT mESCs. ChIP-seq for WT Ezh2, H3K27me3 (WT and EED cage MT) and position of ORCA (<u>O</u>ptical <u>R</u>econstruction of <u>C</u>hromatin <u>A</u>rchitecture) probes is shown below. **b and c**, Violin plots of **(b)** the distance (as measured by ORCA) of indicated regions in WT mESCs and **(c)** the distance of Hmx1 to either Msx1 or Nkx1-1 in WT and Hmx1 deletion mESCs.

**Supplementary Figure 4:**
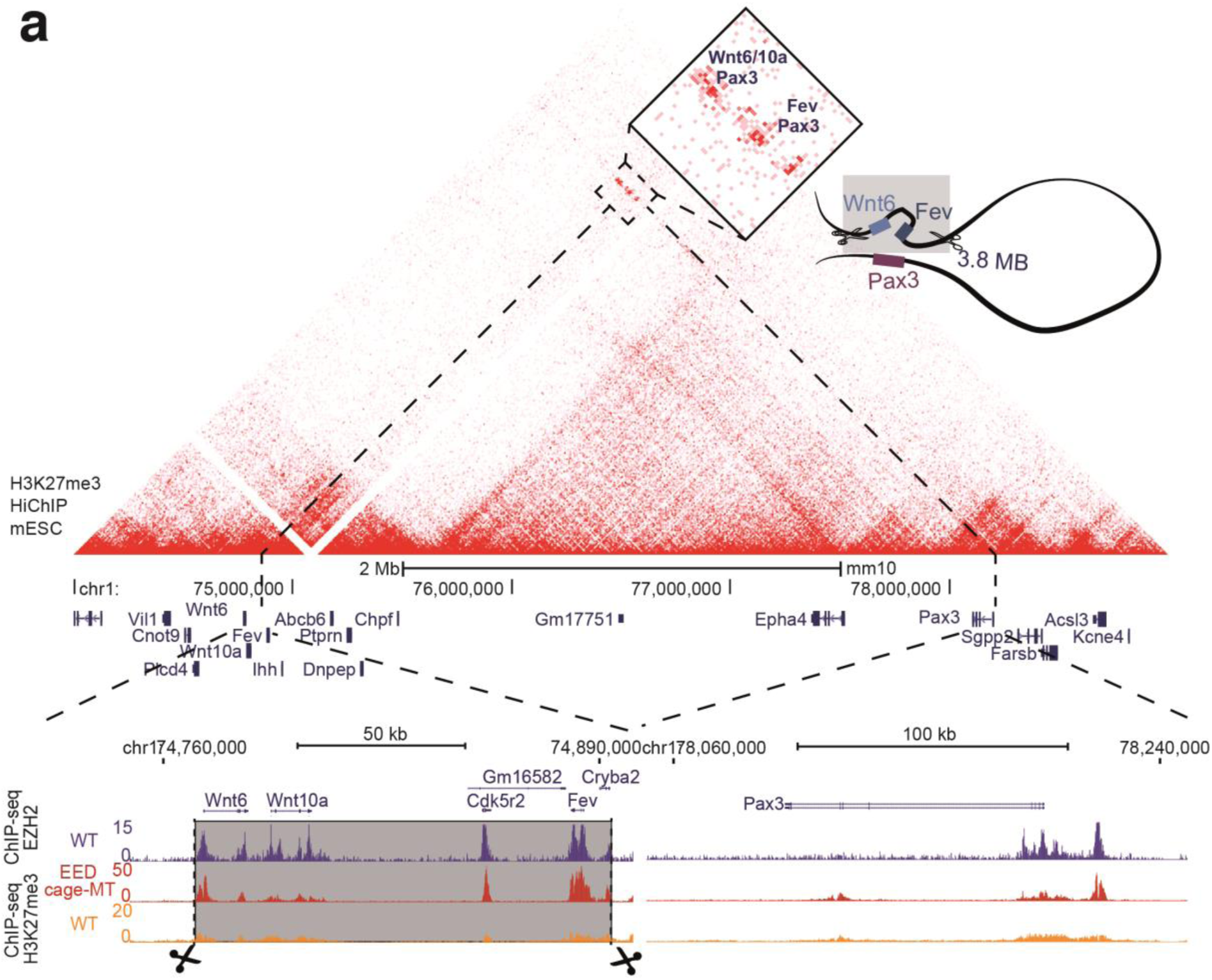
Wnt6 loop anchor deletion. **a**, H3K27me3 HiChIP contact matrix (10 kb resolution) at the Wnt/Pax3 locus in WT mESCs. ChIP-seq for WT EZH2, H3K27me3 (WT and EED cage MT)

**Supplmentary Figure 5:**
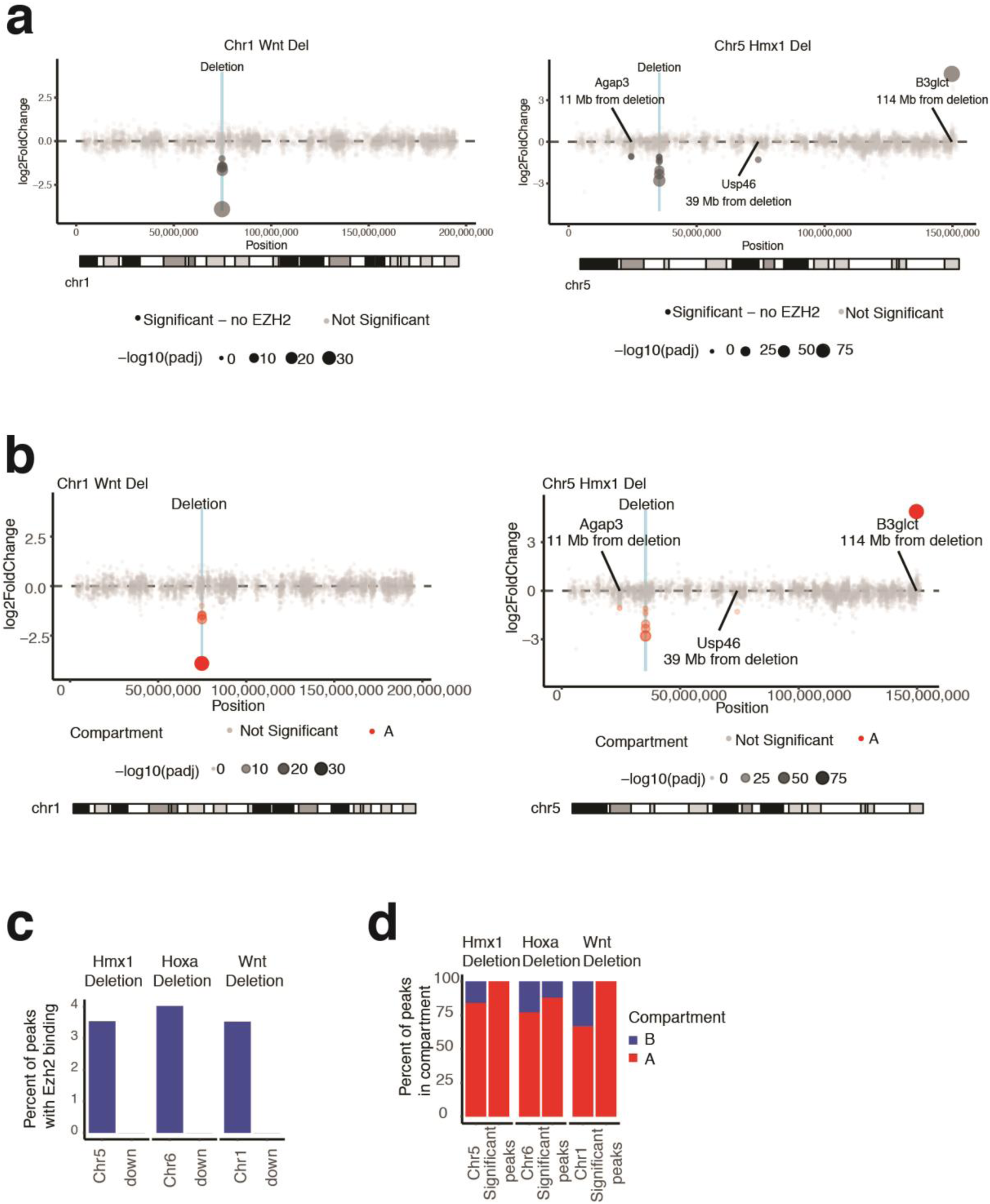
Changes of H3K27me3 signal in anchor point deletions correlated with A/B compartments and EZH2 binding. **a and b**, Scatter plots for the Wnt and Hmx1 anchor point deletions, demonstrating altered H3K27me Cut&Tag signal related to the genomic position on the chromosome and colored by **(a)** binding of EZH2 and **(b)** the type of compartment in which the change occurs. Log2 fold changes and p-values (cutoff of absolute value log2FC > 1 and Benjamini-Hochberg adjusted p-value < 0.05 for significance) calculated in DESeq2 (Love et al ref). **c and d**, Bar plot for the three anchor point deletions describing the percentage of **(c)** significantly downregulated (log2FC < -2, Benjamini-Hochberg adjusted p-value < 0.05) H3K27me3 peaks with EZH2 binding in WT mESCs or **(d)** significantly altered H3K27me3 peaks (absolute value log2FC > 1, Benjamini-Hochberg adjusted p-value < 0.05) colored by A/B compartment in which the H2K27me3 peak is located. Distributions of EZH2 binding and A/B compartments for all H3K27me3 peaks on that chromosome included for comparison.

**Supplementary Figure 6:**
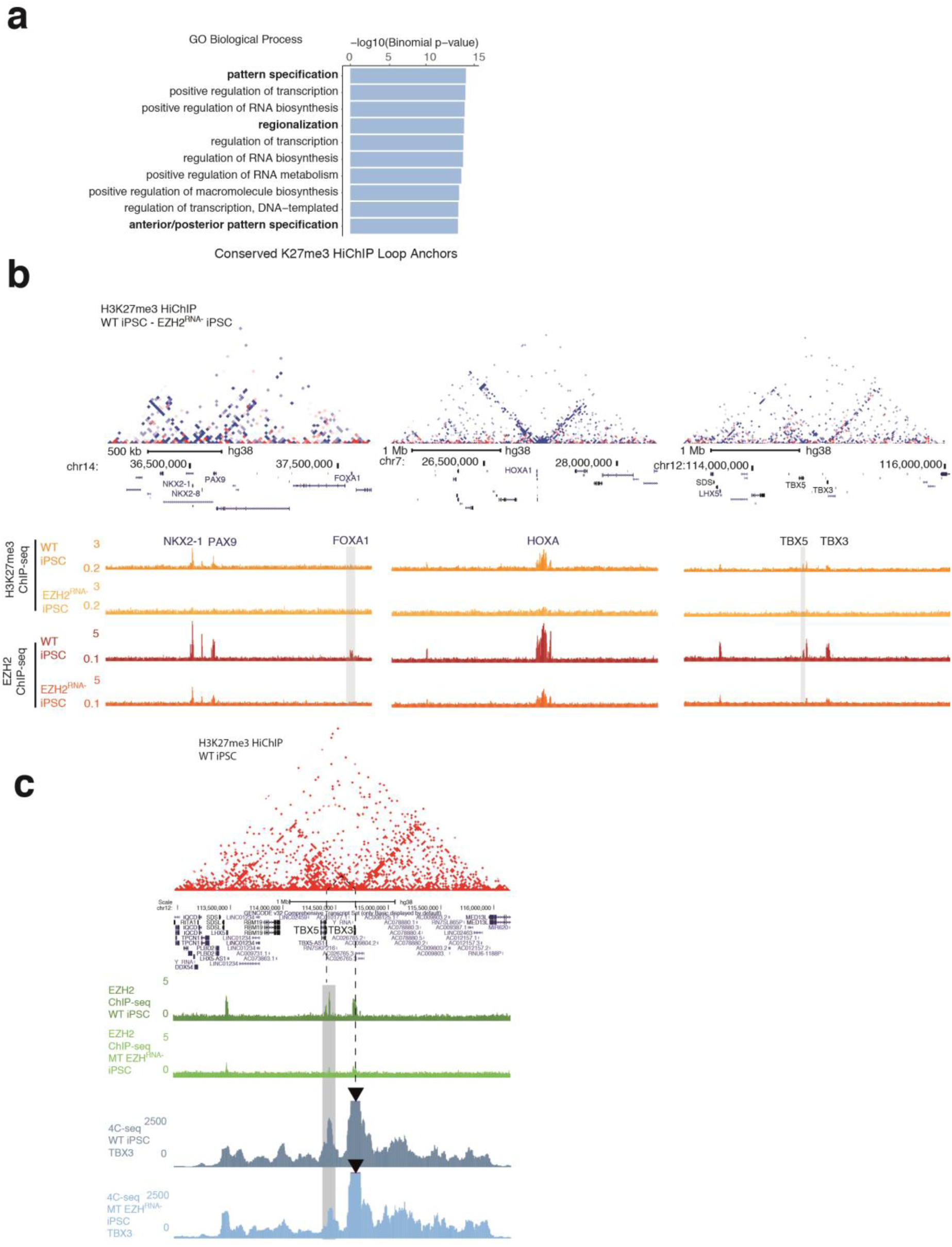
Altered polycomb binding due to loss of RNA binding by EZH2 influences the chromatin conformation and H3K27me3 spreading at different developmental loci. **a**, Gene ontology terms enriched at conserved H3K27me3-associated loop anchors between mESCs and iPSCs. **b**, iPSC H3K27me3 HiChIP subtraction contact matrices (WT – *EZH2*^*RNA-*^, 20 kb resolution) for three representative developmental loci, with corresponding ChIP-seq tracks of H3K27me3 and EZH2 below. Blue color in the subtraction maps corresponds to decreased signal in *EZH2*^*RNA-*^ iPSCs. **c**, H3K27me3 HiChIP contact matrix (20 kb resolution) showing the 3D chromatin configuration at the *TBX3*/*TBX5* locus in WT human iPSCs. EZH2 ChIP-seq and 4C-seq at *TBX3* viewpoint in WT and *EZH2*^*RNA-*^ iPSCs is shown below (n=1. Reduced contact with *TBX5* accompanied by reduced EZH2 binding highlighted.

## Methods

### CRISPR-Cas9-engineered structural variants in mESC

Mouse embryonic stem cells (mESCs) were cultured in ES cell medium containing knockout DMEM 15% FCS and 1000 U/ml LIF. mESC carrying the desired deletions were generated according to the CRISPR-Cas-induced structural variant (CRISVar) protocol_53_. Briefly, per structural variant two single-guide RNAs (sgRNAs) were designed using the “CRISPR guides” design tool of Benchling (https://benchling.com/), picking the guides showing the best off-target score. WT G4 mESCs (129/Sv × C57BL/6 F1 hybrid background)_54_ were co-transfected with the two respective sgRNA-pX459 vectors using the FUGENE HD transfection reagent (Promega) following the manufacturer’s instructions. Individual ESC clones were screened for deletions via PCR, copynumber variation (CNV) qPCR and verified by PCR amplification and Sanger sequencing of the CRISPR-breakpoint. Sequences of sgRNAs, CRISPR breakpoints, genotyping PCR and CNV qPCR primers are listed in Supplementary Table 1.

### Human iPSC

WT an MT iPSC clones where obtained from Thomas R. Cech Lab, MT iPSC clones were generated by Yicheng Long from the original WTC-11 iPSC from Coriell Institute (catalog# GM25256, deposited by Dr. Bruce R. Conklin, Gladstone Institute, UCSF). For regular culture and passaging, cells where maintained in Essential 8 Flex medium (Thermo Fisher A2858501) using Vitronectin (Thermo Fisher A14700) as the coating material and the medium was changed every other day until ready for passaging. To passage the cells, 5-10 min incubation with 0.5 mM EDTA in PBS after a PBS wash step facilitated detachment and cells were split with 1 to 6 (up to 10) ratio into new culture dishes coated with vitronectin. Cells were cryo-preserved in Essential 8 Flex medium with 10% DMSO for long-term storage.

### HiChIP

5×10_6_ cells where fixed in 2% formaldehyde for 10 minutes at room temperature. HiChIP was performed as previously described_12_ using antibodies against H3K27me3 (millipore sigma 07-449) and H3K27ac (am 39133) with the following optimizations_13_ (marked in bold): SDS treatment at 62C for **5 min**; restriction digest for **15 min**; **no heat inactivation** of restriction enzyme, instead wash nuclei twice with 1xrestriction enzyme buffer; biotin fill-in reaction incubation at 37C for **15min**; ligation at room temperature for **2 hours**.

### 4C-Seq

4C-seq libraries were generated from fixed cells as described previously _48_. HindIII (6-bp cutter) was used as primary restriction enzyme. NlaIII was used as secondary restriction enzyme. For each viewpoint, a total of 1.6 mg of each library was amplified by PCR (primer sequence in **Supplementary Table 1**). Samples were sequenced 2×75 bp with Ilumina Hi-Seq 4000 technology according to standard protocols.

### Cut&Tag

Cut&Tag (Cleavage Under Targets and Tagmentation) experiments were performed according to Kaya-Okur et al._45_. In short, cells were harvested with accutase and aliquots of 100.000 cells were conjugated to 10µL of activated Concanavalin A coated beads (Bangs laboratories) per sample. Primary antibody incubation was performed for 2h at RT in 100µL antibody buffer (20 mM HEPES-KOH pH7.5, 150 mM NaCl, 0.5 mM Spermidine, 0.05% Digitonin, 2mM EDTA, 0.1% BSA, 1× potease inhibitors) and either 1µL of H3K27me3 (Active Motif #61017) or IgG (abcam #ab6709) antibody (1:100). Incubation of the secondary antibody was performed for 1h at RT with either rabbit anti-mouse (abcam # ab46540, for H3K27me3) or guinea pig anti-rabbit antibody(Antibodies online #ABIN101961, for IgG) in a 1:100 dilution in Dig-Wash buffer (20 mM HEPES-KOH pH7.5, 150 mM NaCl, 0.5 mM Spermidine, 0.05% Digitonin, 1× potease inhibitors). The pA-Tn5 adapter complex (in-house made batch) was used in a 1:300 dilution in Dig-300 buffer (20 mM HEPES-KOH pH 7.5, 300 mM NaCl, 0.5 mM Spermidine, 0.01% Digitonin, 1× protease inhibitor) and incubated with the samples for 1h at RT. After transposition reaction (1h at 37°C) and reverse-crosslink (overnight at 37°C followed by inactivation of Proteinase K at 70°C for 20min), samples were purified using the Zymo ChIP DNA clean and concentrator kit according to manufacturer’s instructions. PCR was performed with i5/i7 Nextera index primers and NEBNext Hifi 2x PCR mastermix, with a total of 14 cycles. Post-PCR clean-up was carried out by adding 0.9x volume of Ampure XP beads and elution in 20 µL of ultrapure H_2_O.

### ORCA Imaging and data analysis

The primary probes tiling the regions of interest (Supplemental table) were designed as previously described in_14_ with the modification of removing the fiducial labels on primary probes. Separate fiducial probes were designed corresponding to each chromosome of interest (**Supplementary Table 1**) spanning 200kb of the chromosomes tiled by the experimental probes for image registration purposes. Probes were amplified from the oligopool (CustomArray), and amplified according to the protocol described in _14,55_ In preparation for imaging, mES cells were collected and fixed in 4% PFA in 1xPBS for 10 min. Cells were then washed 3x in 1xPBS and stored in 70% ethanol for up to 3 months. 40-mm glass coverslips (Bioptechs) were coated with poly-d-lysine for at least 1 hour and then rinsed with 1xPBS to remove residue. A population of control and deletion mES cells were then plated directly onto the coverslip in two spatially distinct populations and allowed to dry for 7-10 minutes. Once cells were dried and adhered to the slide, the hybridization and imaging were performed as previously described (Mateo et al. 2019). For primary probe hybridization, cells were permeabilized for 10min with 0.5% Triton-X in 1xPBS, the DNA was then denatured by treatment with 0.1M HCL for 5min. 2ug of primary probes in hybridization solution was then added directly on to cells, placed on a heat block for 90C for 3min and incubated overnight at 42C in a humidified chamber. Prior to imaging, the samples were post-fixed for 1h in 8%PFA +2% glutaraldehyde (GA) in 1xPBS. The samples were then washed in 2xSSC and either imaged directly or stored for up to a week in 4C prior to imaging. For imaging, samples were mounted into a Bioptechs flow chamber, and secondary probe hybridization and step by step imaging of individual barcodes, and image processing was performed as in_14_. Image analysis was performed as described in_14_. For all analysis in Figure 2 and supp. Figure 2, ORCA data was processed to calculate absolute distances between all barcodes for all cells. Contact frequency was calculated by calculating the fraction of cells where the probes were within a 200 nm distance. A ‘cell’ refers to all detected spots, with 2 spots per cell, corresponding to each allele. Absolute distances between barcodes were plotted as violin plots to show distribution of distances and wilcoxon rank-sum tests were performed to indicate significant difference between groups. Representative images were created from max projections of cells where contact between regions of interest (within 200nm) was noted.

### HiChIP data processing

HiChIP data were processed as described previously_12_. Briefly, paired end reads were aligned to hg38 or mm10 genomes using the HiC-Pro pipeline_56_ (version 2.11.0). Default settings were used to remove duplicate reads, assign reads to MboI restriction fragments, filter for valid interactions, and generate binned interaction matrices. The Juicer pipeline’s HiCCUPS tool and FitHiChIP were used to identify loops_36,57_. Filtered read pairs from the HiC-Pro pipeline were converted into .hic format files and input into HiCCUPS using default settings. Dangling end, self-circularized, and re-ligation read pairs were merged with valid read pairs to create a 1D signal bed file. FitHiChIP was used to identify “peak-to-all” interactions at 10 kb resolution using peaks called from the one-dimensional HiChIP data. A lower distance threshold of 20 kb was used. Bias correction was performed using coverage specific bias. 1D signal bed files were converted to bigwig format for visualization using deepTools bamCoverage (version 3.3.1) with the following parameters: --bs 5 --smoothLength 105 --normalizeUsing CPM --scaleFactor 10_58_. Enrichment of 1D signal at ChIP-seq peaks was computed using deepTools computeMatrix (version 3.3.1)_58_. TAD and A/B compartment annotations were obtained from a previously published mESC Hi-C dataset_43_. Gene ontology enrichment at loop anchors was performed using GREAT_59_ (version 4.0.4).

### Virtual 4C

Virtual 4C plots were generated from dumped matrices generated with Juicebox. The Juicebox tools dump command was used to extract the chromosome of interest from the HiChIP .hic file or Hi-C .hic file from Bonev *et al*._*43*_ obtained from the 4D Nucleome Data Portal_60_. The interaction profile at the indicated resolution for the bin containing the anchor was then plotted in R following scaling by the total number of filtered reads in each experiment.

### Cut&Tag data processing

Cut&Tag data were processed as described previously_45_ with the following modifications. Paired-end reads were aligned to the mm10 genome using Bowtie2_61_ (version 2.3.4.1) with the --very-sensitive option following adapter trimming with Trimmomatic_62_ (version 0.39). Reads with MAPQ values less than 10 were filtered using samtools and PCR duplicates removed using Picard’s MarkDuplicates. MACS2_62_ (version 2.1.1.20160309) was used for peak calling with the following parameters: macs2 callpeak -t input_bedpe -p 1e-5 -f BEDPE --keep-dup all -g mm -n output_file. A reproducible peak set across biological replicates was defined using the IDR framework (version 2.0.4.2). Reproducible peaks from all samples were then merged to create a union peak set. Cut&Tag signal was converted to bigwig format for visualization using deepTools bamCoverage (version 3.3.1) with the following parameters: --bs 5 --smoothLength 105 --normalizeUsing CPM --scaleFactor 10_58_. Statistical analysis was performed using DESeq2_63_ (version 1.22.2).

### ChIP-seq data processing

ChIP-seq data were obtained from the Gene Expression Omnibus (GEO) using the following accession numbers: mESC EZH2 ChIP-seq_42_ (GSE111433), EED cage mutant mESC H3K27me3 ChIP-seq_6_ (GSE94429), WT and *EZH2*_*RNA-*_ iPSC EZH2 and H3K27me3 ChIP-seq_15_ (GSE128135). Sequencing reads were aligned to the hg38 or mm10 genome using Bowtie2_61_ (version 2.3.4.1) with the --very-sensitive option following adapter trimming with Trimmomatic_64_ (version 0.39). Reads with MAPQ values less than 10 were filtered using samtools and PCR duplicates removed using Picard’s MarkDuplicates. MACS2_64_ (version 2.1.1.20160309) was used for peak calling with the following parameters: macs2 callpeak -t input_bed -f BED -n output_file --nomodel --shift 0 -q 0.01. A reproducible peak set across technical replicates was defined using the IDR framework (version 2.0.4.2). Reproducible peaks from all samples were then merged to create a union peak set. Additional processed ChIP-seq alignments and peak calls were downloaded from ENCODE from the following accessions: mESC H3K27me3 ChIP-seq (ENCSR059MBO), mESC H3K27ac ChIP-seq (ENCSR000CGQ)_35_. ChIP-seq signal was converted to bigwig format for visualization using deepTools bamCoverage (version 3.3.1) with the following parameters: --bs 5 --smoothLength 105 --normalizeUsing CPM --scaleFactor 10_58_. Enrichment of ChIP-seq signal at HiChIP loop anchors was performed using Homer’s annotatePeaks_65_ (version 4.10). Statistical analysis was performed using DESeq2_63_ (version 1.22.2).

### 4C-seq data processing_15_

4C-seq data was processed using pipe4C_66_ and aligned to the hg38 genome using reading primer sequences in **Supplementary Table 1**.

**Supplementary Table 1.**
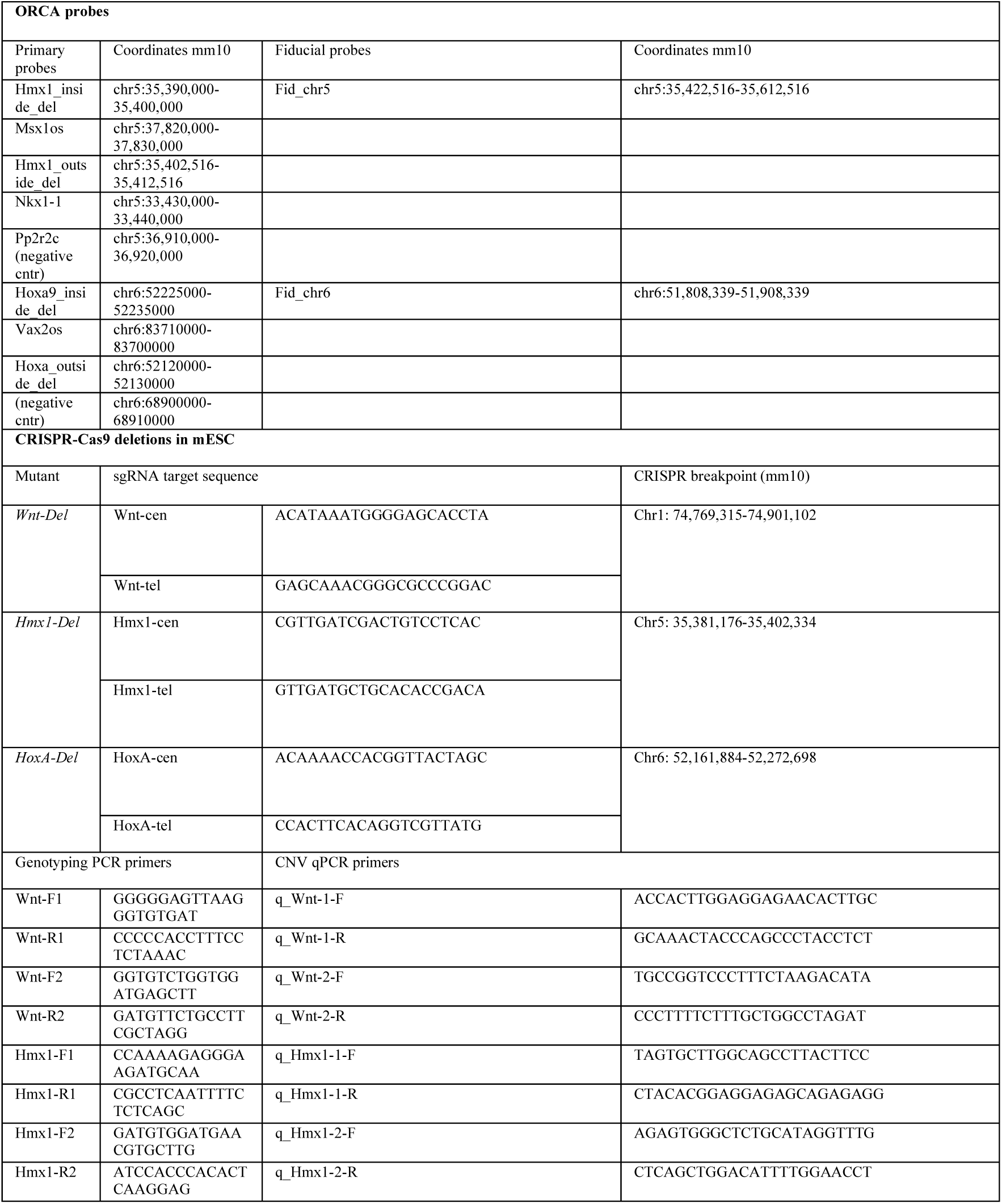
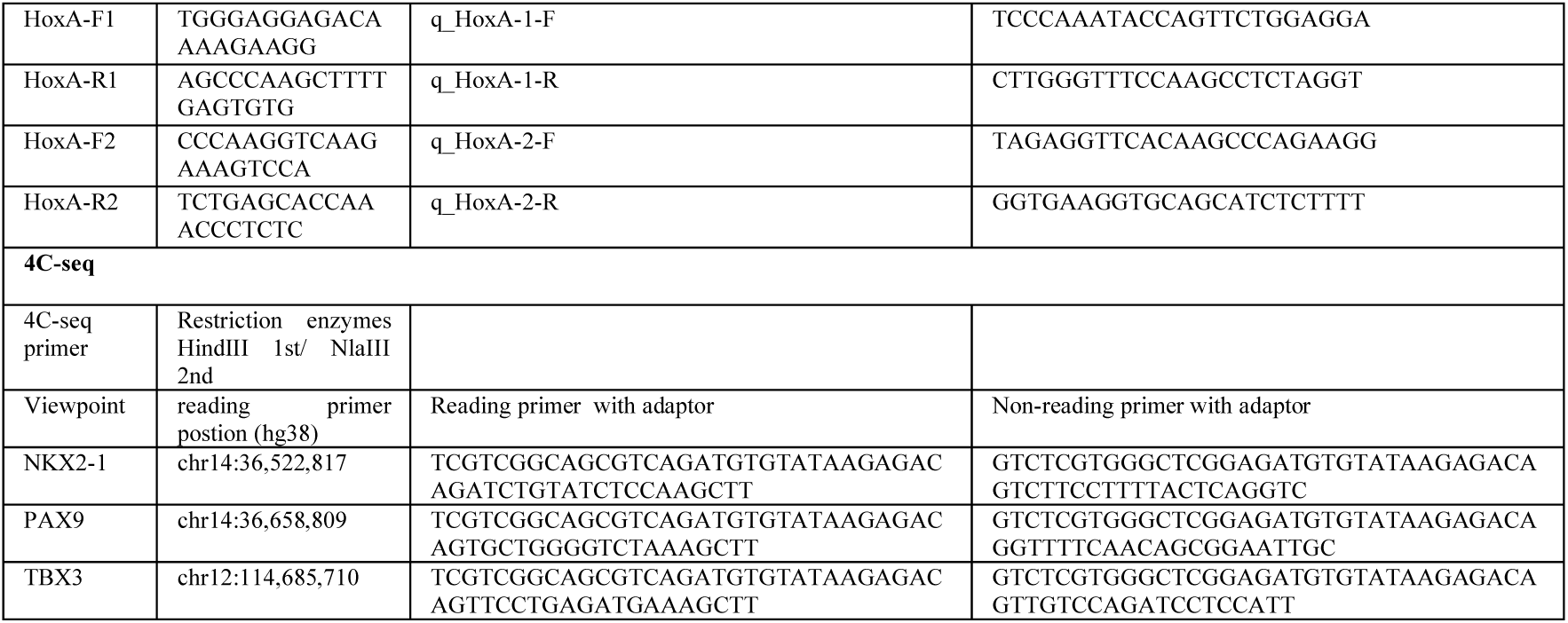

## Reference

1. Schuettengruber, B., Bourbon, H.M., Di Croce, L. & Cavalli, G. Genome Regulation by Polycomb and Trithorax: 70 Years and Counting. Cell 171, 34–57 (2017).

2. Blackledge, N.P., Rose, N.R. & Klose, R.J. Targeting Polycomb systems to regulate gene expression: modifications to a complex story. Nat Rev Mol Cell Biol 16, 643–649 (2015).

3. Boyer, L.A. et al. Polycomb complexes repress developmental regulators in murine embryonic stem cells. Nature 441, 349–53 (2006).

4. Kundu, S. et al. Polycomb Repressive Complex 1 Generates Discrete Compacted Domains that Change during Differentiation. Mol Cell 65, 432–446 e5 (2017).

5. Margueron, R. et al. Ezh1 and Ezh2 maintain repressive chromatin through different mechanisms. Mol Cell 32, 503–18 (2008).

6. Oksuz, O. et al. Capturing the Onset of PRC2-Mediated Repressive Domain Formation. Mol Cell 70, 1149–1162 e5 (2018).

7. Denholtz, M. et al. Long-range chromatin contacts in embryonic stem cells reveal a role for pluripotency factors and polycomb proteins in genome organization. Cell Stem Cell 13, 602–16 (2013).

8. McLaughlin, K. et al. DNA Methylation Directs Polycomb-Dependent 3D Genome Re-organization in Naive Pluripotency. Cell Rep 29, 1974–1985 e6 (2019).

9. Rhodes, J.D.P. et al. Cohesin Disrupts Polycomb-Dependent Chromosome Interactions in Embryonic Stem Cells. Cell Rep 30, 820–835 e10 (2020).

10. Schoenfelder, S. et al. Polycomb repressive complex PRC1 spatially constrains the mouse embryonic stem cell genome. Nat Genet 47, 1179–1186 (2015).

11. Rowley, M.J. et al. Evolutionarily Conserved Principles Predict 3D Chromatin Organization. Mol Cell 67, 837–852 e7 (2017).

12. Mumbach, M.R. et al. HiChIP: efficient and sensitive analysis of protein-directed genome architecture. Nat Methods 13, 919–922 (2016).

13. Mumbach, M.R. et al. HiChIRP reveals RNA-associated chromosome conformation. Nat Methods 16, 489–492 (2019).

14. Mateo, L.J. et al. Visualizing DNA folding and RNA in embryos at single-cell resolution. Nature 568, 49–54 (2019).

15. Long, Y. et al. RNA is essential for PRC2 chromatin occupancy and function in human pluripotent stem cells. Nat Genet (2020).

16. Plath, K. et al. Role of histone H3 lysine 27 methylation in X inactivation. Science 300, 131–5 (2003).

17. Mohammad, F. et al. Kcnq1ot1/Lit1 noncoding RNA mediates transcriptional silencing by targeting to the perinucleolar region. Mol Cell Biol 28, 3713–28 (2008).

18. Okamoto, I., Otte, A.P., Allis, C.D., Reinberg, D. & Heard, E. Epigenetic dynamics of imprinted X inactivation during early mouse development. Science 303, 644–9 (2004).

19. Koppens, M. & van Lohuizen, M. Context-dependent actions of Polycomb repressors in cancer. Oncogene 35, 1341–52 (2016).

20. Deevy, O. & Bracken, A.P. PRC2 functions in development and congenital disorders. Development 146(2019).

21. Schuettengruber, B., Chourrout, D., Vervoort, M., Leblanc, B. & Cavalli, G. Genome regulation by polycomb and trithorax proteins. Cell 128, 735–45 (2007).

22. Mendenhall, E.M. et al. GC-rich sequence elements recruit PRC2 in mammalian ES cells. PLoS Genet 6, e1001244 (2010).

23. Riising, E.M. et al. Gene silencing triggers polycomb repressive complex 2 recruitment to CpG islands genome wide. Mol Cell 55, 347–60 (2014).

24. Joshi, O. et al. Dynamic Reorganization of Extremely Long-Range Promoter-Promoter Interactions between Two States of Pluripotency. Cell Stem Cell 17, 748–757 (2015).

25. Vieux-Rochas, M., Fabre, P.J., Leleu, M., Duboule, D. & Noordermeer, D. Clustering of mammalian Hox genes with other H3K27me3 targets within an active nuclear domain. Proc Natl Acad Sci U S A 112, 4672–7 (2015).

26. Sexton, T. et al. Three-dimensional folding and functional organization principles of the Drosophila genome. Cell 148, 458–72 (2012).

27. Pal, K. et al. Global chromatin conformation differences in the Drosophila dosage compensated chromosome X. Nat Commun 10, 5355 (2019).

28. Ngan, C.Y. et al. Chromatin interaction analyses elucidate the roles of PRC2-bound silencers in mouse development. Nat Genet 52, 264–272 (2020).

29. Xiaotian Zhang, M.J., Xingfan Huang, Jianzhong Su, Muhammad S. Shamim, Ivan D. Bochkov, Jaime Reyes, Haiyoung Jung, Emily Heikamp, Aiden, Wei Li, Erez Lieberman Aiden, Margaret A. Goodell. Large DNA Methylation Nadirs Anchor Chromatin Loops Maintaining Hematopoietic Stem Cell Identity. Molecular Cell 78, P506-521.e6, (2020).

30. Du, Z. et al. Polycomb Group Proteins Regulate Chromatin Architecture in Mouse Oocytes and Early Embryos. Mol Cell 77, 825–839 e7 (2020).

31. Rinn, J.L. et al. Functional demarcation of active and silent chromatin domains in human HOX loci by noncoding RNAs. Cell 129, 1311–23 (2007).

32. Wang, X. et al. Targeting of Polycomb Repressive Complex 2 to RNA by Short Repeats of Consecutive Guanines. Mol Cell 65, 1056–1067 e5 (2017).

33. Bonasio, R. et al. Interactions with RNA direct the Polycomb group protein SCML2 to chromatin where it represses target genes. Elife 3, e02637 (2014).

34. Mumbach, M.R. et al. Enhancer connectome in primary human cells identifies target genes of disease-associated DNA elements. Nat Genet (2017).

35. Yue, F. et al. A comparative encyclopedia of DNA elements in the mouse genome. Nature 515, 355–64 (2014).

36. Rao, S.S. et al. A 3D Map of the Human Genome at Kilobase Resolution Reveals Principles of Chromatin Looping. Cell (2014).

37. Dixon, J.R. et al. Topological domains in mammalian genomes identified by analysis of chromatin interactions. Nature 485, 376–80 (2012).

38. Nora, E.P. et al. Spatial partitioning of the regulatory landscape of the X-inactivation centre. Nature 485, 381–5 (2012).

39. Andrey, G. et al. Characterization of hundreds of regulatory landscapes in developing limbs reveals two regimes of chromatin folding. Genome Res 27, 223–233 (2017).

40. Zhang, X. et al. Large DNA Methylation Nadirs Anchor Chromatin Loops Maintaining Hematopoietic Stem Cell Identity. Mol Cell 78, 506–521 e6 (2020).

41. Boyle, S. et al. A central role for canonical PRC1 in shaping the 3D nuclear landscape. Genes Dev 34, 931–949 (2020).

42. Lee, C.H. et al. Allosteric Activation Dictates PRC2 Activity Independent of Its Recruitment to Chromatin. Mol Cell 70, 422–434 e6 (2018).

43. Bonev, B. et al. Multiscale 3D Genome Rewiring during Mouse Neural Development. Cell 171, 557–572 e24 (2017).

44. Lupianez, D.G. et al. Disruptions of topological chromatin domains cause pathogenic rewiring of gene-enhancer interactions. Cell 161, 1012–1025 (2015).

45. Kaya-Okur, H.S. et al. CUT&Tag for efficient epigenomic profiling of small samples and single cells. Nat Commun 10, 1930 (2019).

46. Christopher M. Weber, A.H., Simon M. G. Braun, Alistair N. Boettiger, Gerald R. Crabtree. bioRxiv (2020).

47. Long, Y. et al. Conserved RNA-binding specificity of polycomb repressive complex 2 is achieved by dispersed amino acid patches in EZH2. Elife 6(2017).

48. van de Werken, H.J. et al. 4C technology: protocols and data analysis. Methods Enzymol 513, 89–112 (2012).

49. Elgin, S.C. & Reuter, G. Position-effect variegation, heterochromatin formation, and gene silencing in Drosophila. Cold Spring Harb Perspect Biol 5, a017780 (2013).

50. Khalil, A.M. et al. Many human large intergenic noncoding RNAs associate with chromatin-modifying complexes and affect gene expression. Proc Natl Acad Sci U S A 106, 11667–72 (2009).

51. Jitendra Thakur, H.F., Trizia Llagas, Christine M. Disteche, Steven Henikoff. Architectural RNA is required for heterochromatin organization. bioRxiv (2019).

52. Amaral, P.P. et al. Genomic positional conservation identifies topological anchor point RNAs linked to developmental loci. Genome Biol 19, 32 (2018).

53. Kraft, K. et al. Deletions, Inversions, Duplications: Engineering of Structural Variants using CRISPR/Cas in Mice. Cell reports (2015).

54. George, S.H. et al. Developmental and adult phenotyping directly from mutant embryonic stem cells. Proc Natl Acad Sci U S A 104, 4455–60 (2007).

55. Boettiger, A.N. et al. Super-resolution imaging reveals distinct chromatin folding for different epigenetic states. Nature 529, 418–22 (2016).

56. Servant, N. et al. HiC-Pro: an optimized and flexible pipeline for Hi-C data processing. Genome Biol 16, 259 (2015).

57. Bhattacharyya, S., Chandra, V., Vijayanand, P. & Ay, F. Identification of significant chromatin contacts from HiChIP data by FitHiChIP. Nat Commun 10, 4221 (2019).

58. Ramirez, F. et al. deepTools2: a next generation web server for deep-sequencing data analysis. Nucleic Acids Res 44, W160–5 (2016).

59. McLean, C.Y. et al. GREAT improves functional interpretation of cis-regulatory regions. Nat Biotechnol 28, 495–501 (2010).

60. Bonev, B. & Cavalli, G. Organization and function of the 3D genome. Nat Rev Genet 17, 772 (2016).

61. Langmead, B. & Salzberg, S.L. Fast gapped-read alignment with Bowtie 2. Nat Methods 9, 357–9 (2012).

62. Zhang, Y. et al. Model-based analysis of ChIP-Seq (MACS). Genome Biol 9, R137 (2008).

63. Love, M.I., Huber, W. & Anders, S. Moderated estimation of fold change and dispersion for RNA-seq data with DESeq2. Genome Biol 15, 550 (2014).

64. Bolger, A.M., Lohse, M. & Usadel, B. Trimmomatic: a flexible trimmer for Illumina sequence data. Bioinformatics 30, 2114–20 (2014).

65. Heinz, S. et al. Simple combinations of lineage-determining transcription factors prime cis-regulatory elements required for macrophage and B cell identities. Mol Cell 38, 576–89 (2010).

66. Krijger, P.H.L., Geeven, G., Bianchi, V., Hilvering, C.R.E. & de Laat, W. 4C-seq from beginning to end: A detailed protocol for sample preparation and data analysis. Methods 170, 17–32 (2020).

